# Do Neotropical adult dobsonflies (Megaloptera, Corydalidae) feed on solid material in the nature? Information from a gut content study

**DOI:** 10.1101/2022.04.19.488733

**Authors:** Hugo Alejandro Álvarez, José Manuel Tierno de Figueroa, Jorge Alejandro Cebada-Ruiz

## Abstract

Females of two species of corydalids (Megaloptera), *Corydalus magnus* Contreras-Ramos, 1998 and *Platyneuromus soror* (Hagen, 1861), are studied with the aim to determine if solid food is present in their guts, which would indicate the existence of feeding on solid food during the imaginal life in nature. Gut anatomical architecture is also studied and described for females of both species, showing no peculiarities regarding the existing information in other close related species and confirming that digestive morphology is conserved in Megaloptera, as previously reported in literature. No significant solid food was detected in any of the species, so in the nature, they may rely more on liquid sources and/or fat reserves produced in the larval stage for the energetic requirements of adult life.

## Introduction

The natural habits of adult megalopterans remain largely unexplored. The order Megaloptera is composed of holometabolous insects, with aquatic larvae and terrestrial pupae and adults (Cover and Bogan 2015). This order is composed by two families, Sialidae (alderflies) and Corydalidae, the latter including the subfamilies Corydalinae (dobsonflies) and Chauliodinae (fishflies). The life cycle duration of most species ranges from one to two years, though some species live up to five years (Elliott 1996; Cover and Resh 2008; Álvarez 2012). Nevertheless, the adult lifespan is short, only from one week to one month (Hayashi 1993; Álvarez 2012, 2014; Cover and Bogan 2015; Villagomez and Contreras-Ramos 2017). Adult dobsonflies are typically nocturnal (Álvarez 2014; Álvarez et al. 2017a; 2019), while many alderflies and fishflies are diurnal (Stange 1990; Elliott 1996; Anderson 2009).

Regarding the feeding habits of adults in nature, in diurnal alderflies, some species are known to visit flowers for nectar or pollen (Knuth 1906; Kato 2000; Takeuchi 2008). In nocturnal dobsonflies adults are presumed to feed on liquid solutions and tree sap (Champlain and Kirk 1926; Parfin 1952; Ishizawa 1988), whereas in nocturnal fishflies adults of few species forage on the flowers for pollen and nectar (Sugiura and Miyazaki 2021) or on tree sap and small insects (Yoshida et al. 1985). However, this information is based on just a few confirmed records of the use of natural food sources in adult megalopterans and data on adult feeding in this group remain limited and sometimes contradictory. So, although some studies pointed out that different *Sialis* species (alderflies) feed on pollen and nectar, no such content had been found in their guts according to Elliott (1996). Afterwards, Tierno de Figueroa and Palomino-Morales (2002) conducted a study analysing the gut content of an adult alderfly species, *Sialis nigripes* Pictet, 1865, and found a very small quantity of debris (that could have being accidentally ingested while drinking) in 2 of the 22 studied males, indicating that males probably do not feed on solid matter in their adult stage. In that study, the nine studied females had, together with a nonsignificant quantity of spores, detritus and pollen grains, rests of the male’s spermatophore inside their guts, indicating that females could obtain considerable additional energy from this source. The observation of females of this species bending their abdomens to reach the spermatophore time after mating to actively ingest the male’s spermatophore supported this hypothesis (Tierno de Figueroa and Palomino-Morales 2002) and coincides with what was previously reported in other Megaloptera, such as *Protohermes* spp. (Hayashi 1992, 1993). So, it is though that, at least some adult megalopterans do not feed on solid materials and depend only on larval reserves and/or liquid food, while other species, or only females of some of those species ingesting the male spermatophores, feed on solid matter during their imaginal life.

Recently, it has been suggested the capability of Neotropical adult dobsonflies to feed on solid fruits under laboratory conditions (Villagomez and Contreras-Ramos 2017). In their study Villagomez and Contreras-Ramos (2017) described the acceptance of different solid foods by individuals of three species of dobsonflies under laboratory conditions and showed that some species were capable of only licking fluids from fruits and other species chewed and swallowed small solid pieces of fruits. Although such laboratory evidence was not a reproduction of natural conditions, it appears to be important and leads to some questions on adult feeding habits, for example, amongst New World species of Corydalinae, is the feeding on solid food a common feature in nature, as showed in some other species of Megaloptera previously studied? If so, is the anatomy of the gut and the ingested food diverse amongst genera and species? Related to this, the aim of the present study is to investigate the possible ingestion of solid food in nature in two species of New World dobsonflies. For that, we describe the gut of females in two species, *Corydalus magnus* Contreras-Ramos, 1998 and *Platyneuromus soror* (Hagen, 1861), and then we examine the gut content by tissue clearing.

## Material and methods

### Research organisms

The studied species, *Platyneuromus soror* (Figure 1a), is characterised by a small to medium body size, thin antennae, a pale-yellow colour with lateral margin of prothorax, brown postocular flanges, and pale brown medium size spots on the wings (Glorioso 1981; Glorioso and Flint 1984).The other studied species, *Corydalus magnus* (Figure 1b), is characterised by a medium to very large body size, very long antennae (sinuate or subserrate), a pallid brown colour with big moderately dark spots at the edge of the wings, small white spots on the wings, and very slightly to moderately patterned heads (Contreras-Ramos 1998).

**Figure 1.**
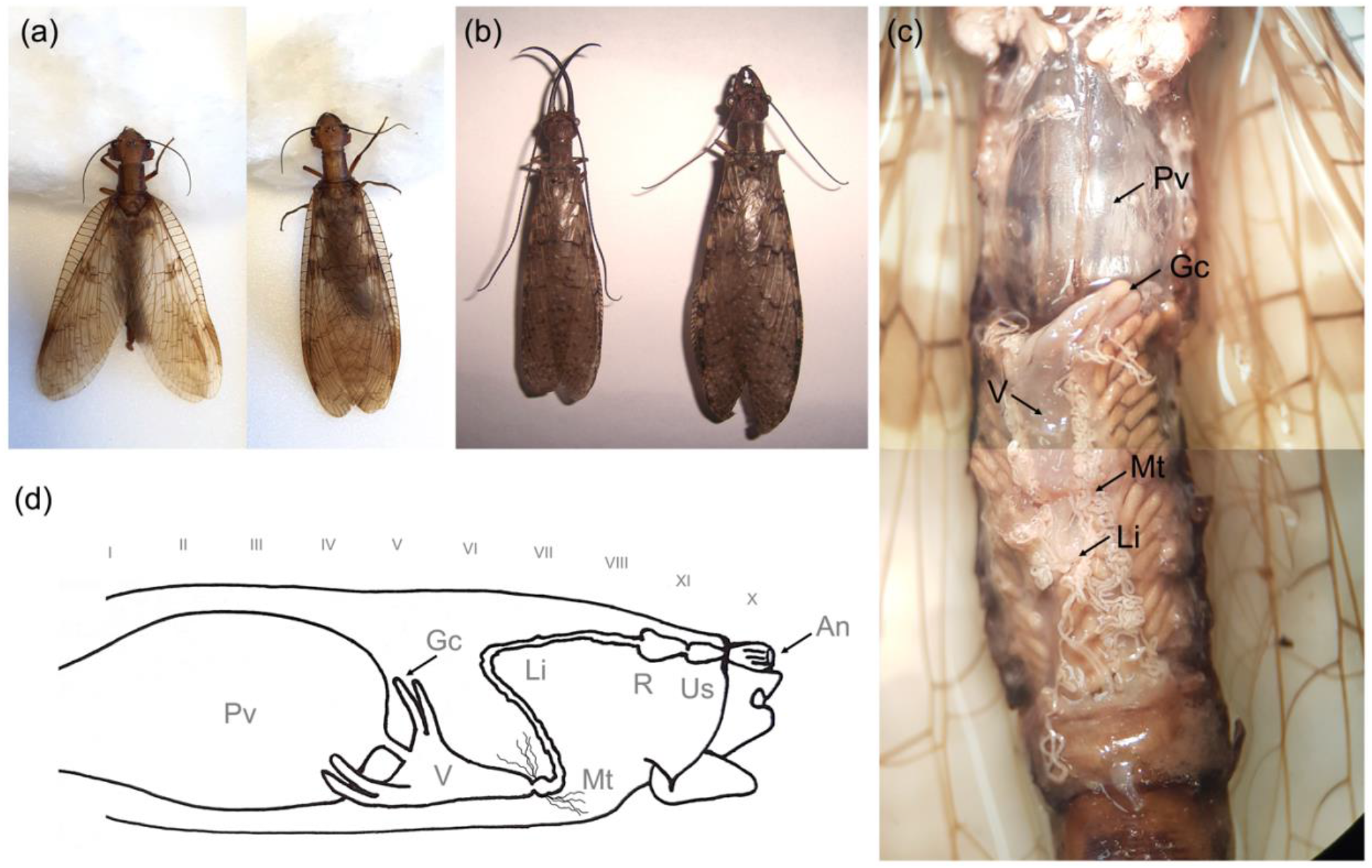
Dobsonflies *Platyneuromus soror* (a) and *Corydalus magnus* (b): male and female habitus (left and right, respectively). Ventral view of a female’s abdomen internal morphology (c) showing the ventriculus (V), gastric caeca (Gc), Malpighian tubules (Mt), and a section of the large intestine (Li) between and surrounded by the ovaries that reach the air filled proventriculus (Pv). Lateral scheme of a female’s gut morphology (d) showing how is positioned the gut in the abdomen, the ventral to dorsal turn of the large intestine, and the position of the rectum (R), urinary sac (Us), and anus (An).

### Specimen collection

Following Álvarez et al. (2019), adult specimens were collected near the streetlights located along the pedestrian paths of the residual water treatment plant of the city of Zacatlán (SOSAPAZ), located in the municipality of Zacatlán (19°56’08.42’N, 97°57’38.21”W), in the Northern Sierra of Puebla, Mexico. During the rainy season of 2018 (late spring), adults were collected at night using entomological aerial nets. Specimens were individually placed in plastic containers with a 70 % alcohol solution and transported to the Laboratory of BioSciences at the faculty of Medicine, Autonomous University of Puebla, Mexico. Then, in the summer of 2019 specimens were sent to the Department of Zoology, University of Granada, Spain, for their dissection and analysis.

### Gut content study

Twelve females of *C. magnus* and nine females of *P. soror* were dissected and their guts were extracted for study their possible content. To extract the complete gut without harming it, adults were dissected with a bistoury through the frontal plane separating the frontal and dorsal sections of the thorax and the abdomen reaching the 10 tergite and sternite, next the gut was pull out of the head. Finally, the end of the abdomen was separated from the dissected sections to conserve the annus with all the gut. Each gut was placed in a vial with Hertwig’s liquid at environmental temperature (approximately 25°C) for approximately 20 hours. Afterwards, guts were cleaned of the adhered remains of the reproductive system and fat body with the help of tweezers and mounted (also in Hertwig’s liquid) on glass slides with cover slips for microscopic examination. A compound microscope Zeiss Primo Star© equipped with ocular micrometer was used, if needed, to estimate the % absolute gut content (% of the whole digestive tract occupied by content) at 40× magnification and the relative abundance of food items in the gut content (% of the total gut content occupied by each component) at 400×. The employed methodology was a modification of that proposed by Bello and Cabrera (1999), previously used in gut content studies of both adults and nymphs/larvae of aquatic insect orders (e.g., Tierno de Figueroa and Sánchez-Ortega 1999; López-Rodríguez et al. 2009; Tierno de Figueroa et al. 2019), including also Megaloptera belonging to the family Sialidae (Zamora-Muñoz et al. 1999; Tierno de Figueroa and Palomino Morales 2002).

## Results

### Morphology of the gut

When dissected, females of both species showed the same anatomical architecture in the gut. The foregut is formed by a tin elongated oesophagus that, within the abdomen, dilatates into a distended air filled proventriculus, which fills the anterior section of the abdomen reaching the ovaries and ovarian tubules approximately at the fifth abdominal segment (Figure 1c). In the midgut the ventriculus joints the proventriculus by a narrow section that bends to the ventral part of the abdomen. The ventriculus, which is contracted and fusiform with two pairs of tubular and obtuse gastric coeca at its sides, is placed at the ventral part of the abdomen surrounded by the ovarian tubules (Figure 1d). In the hindgut, the ventriculus abruptly contracts in an ileum-resembling section giving place to several long and convoluted Malpighian tubules and to the large intestine. Then the large intestine bends to the dorsal part of the abdomen passing by the middle between the ovaries, which is compressed by the development of egg. The lower third of the large intestine is formed by a fusiform coecum that opens into the rectum, which proceeds as a canal fused with a urinary sack to the anus (Figure 1d).

### Gut content

Only in three females of *Corydalus magnus* some negligible solid material was detected in their guts consisting on a few fungi spores and detritus (never occupying more than a 1% of the gut content). Two *Platyneuromus soror* females had one pollen grain each one in their guts that, as in the case of *C. magnus* did not occupy even 1% of the gut. Whit this technique it is not possible (or highly difficult) to discriminate the liquid contents, but we recognise that there were liquids in the gut.

## Discussion

The observations reported here seem to indicate that Neotropical adult dobsonflies do not feed on solid food in the nature. At least for the adult females of the two species studied here, *Corydalus magnus* and *Platyneuromus soror,* more than the 99 % of the gut is absent of solid materials and the low quantity of solid food present could have been ingested accidentally while drinking. In fact, it has been recorded worldwide that adults of several species of Megaloptera may ingest liquids (reviewed by Sugiura and Miyazaki 2021; Villagomez and Contreras-Ramos 2017), and our analysis supports this statement for the studied dobsonflies. Particularly, at least some species of small or mean-sized megalopterans may need to ingest solid food to produce spermatophore in males or to mature eggs in females, as it is the case of the Asian fishfly *Neochauliodes amamioshimanus,* that feeds on the pollen and nectar of *S. wallichii* ssp. *noronhae* flowers (Sugiura and Miyazaki 2021). However, for the case of Neotropical adult dobsonflies, the production of spermatophores has not been recorded, but it is known that adult females of some Nearctic dobsonflies, as *Corydalus cornutus* (Limnaeus, 1758), must feed to provide energy to yolk eggs, produce cases and for maintenance of reproductive activities during their short adult life (Brown and Fitzpatrick 1978). Particularly the here studied great-sized dobsonfly species may rely more on liquid sources and/or fat reserves produced in the larval stage. This fact has been reported in other aquatic insects as Plecoptera, in which only the great sized species do not ingest solid food, or only a non-significant quantity of it, during the adult life (Tierno de Figueroa and López-Rodríguez 2019)

Regarding the capability of Neotropical adult dobsonflies to eat solid material, it is undeniable that some species may be able to ingest solid food (Villagomez and Contreras-Ramos 2017). However, the ingestion of solid food in laboratory conditions does not mean to be necessarily a rule in nature. As Villagomez and Contreras-Ramos (2017) showed, dobsonflies were capable to chew some fruits only when the fruit pulp was exposed, otherwise dobsonflies did not do it. This behaviour suggests that dobsonflies may be attracted to sweet compounds, so they can exploit the situation where the pulp of a fruit is exposed and drink the liquids provided by it, chewing the pulp if needed. Indeed, some studies and several entomologists have reported that Neotropical adult dobsonflies had been collected on traps with fruit baits (Champlain and Kirk 1926; Parfin 1952; personal observations). This could explain the existence of few debris within the gut of the species studied here, which will be an accidental ingestion. In the same manner, the few pollen grains or fungi spores detected could be accidentally ingested by females while drinking on nectar or tree sap, as reported above.

On the other hand, our study describes the morphology of the gut in these two species of dobsonflies. The gut is almost the same in the two species, and it follows the morphology described 150 years ago by Leidy (1849) for the Nearctic *Corydalus cornutus*. Thus, the idea that the anatomy of megalopterans is conserved (Glorioso 1981) is verified here. Moreover, it is outstanding the enlarged and air filled proventriculus. We want to emphasise this feature, because it may be a necessary function for the movement of the soft abdomen in mating, as it has been described in several species of megalopterans, in which the abdomen is completely bend or curled by males (e.g., Álvarez et al. 2017a, 2019; Tierno de Figueroa and Palomino-Morales 2002; Contreras-Ramos 1999) and females (e.g., Hayashi 1993, Tierno de Figueroa and Palomino-Morales 2002). Indeed, in a similar preliminary study of the anatomy of a Neotropical dobsonfly (Álvarez et al. 2017b), in which freshly slaughtered adults were dissected, the abdomen maintained for few minutes their movements and it was observed that the proventriculus was expanding and contracting (personal observation). So, it is possible that an empty proventriculus may enable extreme positions and movements to produce a successful mating, however this remains unclear.

Finally, although there are some reports of the feeding habits of Neotropical adult dobsonflies, the present study supports the absence of solid food in natural conditions within the gut of dobsonflies, and thereby it contributes to broadening our understanding of nocturnal corydalids trophic ecology. Future research on resource allocation acquired during the larval stage, maturation process, etc., would allow a better understanding on the adult biology of Neotropical dobsonflies. In addition, further anatomical studies and comparisons are needed to truly understand external features (but see Liu et al. 2015) and behaviour patterns in the order Megaloptera.

## Acknowledgements

The authors thank the directors and working personnel of ‘Planta de Tratamiento de Aguas Residuales SOSAPAZ’ for providing the facilities for fieldwork, Marisol Rodríguez for her help during fieldwork, and Manuel J. López-Rodríguez for his valuable comments on the manuscript.

